# Schizophrenia hiPSC neurons display expression changes that are enriched for disease risk variants and a blunted activity-dependent response

**DOI:** 10.1101/062885

**Authors:** Panos Roussos, Boris Guennewig, Dominik C. Kaczorowski, Guy Barry, Kristen J. Brennand

## Abstract

**IMPORTANCE:** Schizophrenia (SCZ) is a common illness with complex genetic architecture where both common genetic variation and rare mutations have been implicated. SCZ candidate genes participate in common molecular pathways that are regulated by activity-dependent changes in neurons, including the signaling network that modulates synaptic strength and the network of genes that are targets of fragile X mental retardation protein. One important next step is to further our understanding on the role of activity-dependent changes of genes expression in the etiopathogenesis of SCZ.

**OBJECTIVE:** To examine whether neuronal activity-dependent changes of gene expression is dysregulated in SCZ.

**DESIGN, SETTING, AND PARTICIPANTS:** Neurons differentiated from human induced pluripotent stem cells (hiPSCs) derived from 4 cases with SCZ and 4 unaffected controls were depolarized using potassium chloride. RNA was extracted followed by genome-wide profiling of the transcriptome.

**MAIN OUTCOMES AND MEASURES:** We performed differential expression analysis and gene co-expression analysis to identify activity-dependent or disease-specific changes of the transcriptome. Further, we used gene set analyses to identify co-expressed modules that are enriched for SCZ risk genes.

**RESULTS:** We identified 1,669 genes that are significantly different in SCZ-associated vs. control hiPSC-derived neurons and 1,199 genes that are altered in these cells in response to depolarization. We show that the effect of activity-dependent changes of gene expression in SCZ-associated neurons is attenuated compared to controls. Furthermore, these differentially expressed genes are co-expressed in modules that are highly enriched for genes affected by genetic risk variants in SCZ and other neurodevelopmental disorders.

**CONCLUSIONS AND RELEVANCE:** Our results show that SCZ candidate genes converge to gene networks that are associated with a blunted effect of activity-dependent changes of gene expression in SCZ-associated neurons. Overall, these findings show that hiPSC neurons demonstrate activity-dependent transcriptional changes that can be utilized to examine underlying mechanisms and therapeutic interventions related to SCZ.

## INTRODUCTION

Recent genetic studies have implicated numerous common and rare variants in risk for schizophrenia (SCZ)^1–7^. So far, we have learned that the genetic architecture for SCZ is highly polygenic and will necessitate the study of multiple genes. One of the next challenges is to further understand the biological mechanisms of the large number, and diversity, of genes that are associated with SCZ. To do that, we need to generate functional data capturing molecular pathways that are relevant to SCZ and examine the involvement of multiple candidate genes. Such strategy holds the potential to advance our understanding of the underlying molecular basis of SCZ, as well as a way to develop novel treatments.

Recent studies suggest that a convergent molecular pathway dysregulated in SCZ is the signaling network that modulates synaptic strength or the network of genes that are targets of fragile X mental retardation protein (FMRP)^2,7^. In addition, these networks are enriched for genes affected by mutations in autism and intellectual disability^7^, providing further insight for a pathophysiology shared among neurodevelopmental disorders. Interestingly, both gene networks are composed of many proteins for which expression or function is regulated by neuronal activity^8^, further suggesting that SCZ might be driven by dysregulation of neuronal activity-dependent synapse development and function.

Neurons differentiated from human-induced pluripotent stem cells (hiPSC) can assess human brain cellular properties^9^ and have the potential to detect changes of gene expression in response to neuronal depolarization. We used hiPSC neurons derived from patients with SCZ and unaffected controls and identified disease-specific differences in gene expression as well as blunted effect of activity-dependent changes of gene expression in SCZ compared to controls. Furthermore, these differentially expressed genes are co-expressed in modules that are highly enriched for genes affected by genetic risk variants in SCZ and other neurodevelopmental disorders. Overall, these findings show that SCZ candidate genes converge to gene networks that are associated with differential effect of activity-dependent changes of gene expression in SCZ.

## METHODS

### Generation of hiPSC neurons and RNA-seq

Forebrain neural progenitor cells (NPCs) previously differentiated from case and controls hiPSCs as reported^9,10^ were subjected to 6-weeks of neuronal differentiation and maturation (see **eMaterial** in the Supplement) in 6-well format. Neurons were treated with 50nM KCl (or PBS vehicle control) for the final three hours prior to harvest. Total RNA was isolated from approximately density-matched plates (initially seeded with ~200,000 NPCs per well) using TRIzol (Thermo Fisher), according to the manufacturer’s instructions. The RNA Integrity Number (RIN) was determined using an RNA Nano chip (Agilent Technologies) on the Agilent 2100 Bioanalyzer. All samples have high RIN (mean ± standard deviation: 9.53 ± 0.22). 500 ng of total RNA was used as input material for library preparation using the TruSeq Stranded Total RNA Sample Prep Kit (LT) (Illumina, USA) according to manufacturer’s instructions. Individual libraries were indexed as recommended by Illumina.

### Preprocessing of RNA-seq data and differential expression analysis

Reads were mapped to hg19 reference genome using TopHat (version 2.0.9) and Bowtie (version 2.1.0). Known Ensembl gene levels (version 70) were quantified by HTSeq (version 0.6.0). After exploratory analysis (see **eMaterial** in the Supplement) we identified and included the following five covariates in the differential gene expression analysis: Diagnosis (control and SCZ), Treatment (PBS and KCl), Gender (Male and female), Age and RIN. We used the *voomWithQualityWeights* function from the *limma* package to model the normalized read counts. For each transcript, we fit weighted least-squares linear regression models for the effect on gene expression of each variable on the right-hand side, using linear regression utilities in the limma package:

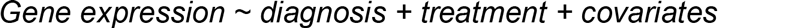

P-values were then adjusted for multiple hypotheses testing using false discovery rate (FDR) estimation with Benjamini and Hochberg correction^11^, and the differentially expressed genes were determined as those with an estimated FDR ≤ 0.05.

### Weighted gene co-expression analysis (WGCNA)

We constructed unsigned gene co-expression networks using the WCGNA package in R, starting with the normalized and residualized (removing effect of age, gender and RIN) expression data for 13,903 genes. For the current data, we used an *R*^2^ cutoff of 0.85, which corresponded to a selection of (² = 7. The minimum module size was set to 30 genes and the minimum height for merging modules was set at 0.3. Ordered from largest (the module containing the most genes) to smallest, each module is sequentially assigned a color name. The less well-connected genes are arbitrarily grouped in the gray module.

### Gene sets for enrichment analyses

To further characterize the DEGs and co-expression genes, we performed enrichment analysis, using a group of gene sets for known molecular pathways and biological processes, including: Gene Ontology (GO) sets of molecular functions (MF), biological processes (BP), and cellular components (CC) (http://www.geneontology.org)^12^; the Reactome dataset (http://www.reactome.org)^13^; and the HUGO Gene Nomenclature Committee (HGNC) gene families (http://www.genenames.org)^14^. In addition, to further characterize the data-driven co-expressed modules, we generated a group of gene sets derived from previous SCZ genetic findings, cell type-specific studies or co-expression analyses (see **eMaterial** in the Supplement). The genes in each module were tested for overlap using Fisher’s exact test and FDR correction across all modules and all gene sets tested.

### Enrichment of modules with genetic risk loci

We tested the enrichment of modules for genes found in genetic loci previously associated with SCZ, including: a) 108 loci discovered in a common variant genome-wide association (GWAS) metaanalysis study of 36,989 SCZ cases and 113,075 controls^1^, b) a literature consensus of 12 copy number variant (CNV) regions collated from numerous rare CNV studies^15^, and c) 756 nonsynonymous (mostly missense, but also including 114 loss-of-function [nonsense, essential splice site, or frameshifting indels]) *de novo* mutations discovered from exome-sequencing across 1,024 SCZ probands and their parents^2–6^. In addition, we used published *de novo* mutations across three neurodevelopmental disorders, including: a) autism spectrum disorders (ASD): 3,446 nonsynonymous and 579 loss-of-function mutations^16–19^, b) intellectual disability (ID): 259 nonsynonymous and 67 loss-of-function mutations^20–23^, and c) epilepsy: 341 nonsynonymous and 58 loss-of-function mutations^24,25^. See **eMaterial** in the Supplement for more details in the enrichment analysis.

## RESULTS

### Differential expression between cases with SCZ and controls

In previous publications, we directly reprogrammed fibroblasts from four SCZ patients and six controls into hiPSCs and differentiated these disorder-specific hiPSCs into forebrain neural progenitor cells (NPCs)^10^ and neurons^9^. As previously reported, spatial patterning, replication and propensity towards neuronal differentiation of NPCs does not appear to vary based on psychiatric diagnosis^9,10^. When differentiated to neurons, forebrain NPCs yield a population of forebrain hiPSC neurons that are VGLUT1-positive, and so are presumably excitatory glutamatergic neurons, although approximately 30– of neurons are GAD67-positive (GABAergic)^9^.

In this study we compared global transcription of hiPSC forebrain neurons from four control subjects and four patients with SCZ (**eTable 1 in the Supplement**), with or without potassium chloride-induced depolarization (PCID). RNA sequencing was performed and we obtain, on average, 21.5 million paired-end reads, among which 94% were mapped on the genome (**eTable 1 in the Supplement**). Following data normalization, there were 13,903 genes for analysis of which 12,678 were protein coding. Multidimensional scaling separated control from SCZ and PBS from KCl-treated samples (**Figure 1A**). We first explored transcriptional changes related to SCZ by identifying differentially expressed genes (DEGs) between SCZ and control neuronal samples. We identified 1,669 DEGs at false discovery rate (FDR) ≤ 0.05 (**eTable 2 in the Supplement**), as illustrated by the heat map (**Figure 1B**) and volcano (**Figure 1C**) plots. Among the 1,669 DEGs, 854 were up-regulated and 815 were down-regulated in SCZ, with a moderate to strong effect (average log2 fold change 3.91; range 0.77-14.50). **Figure 1D** shows illustrative examples for two DEGs that have also been previously associated with SCZ genetic loci^1^: *CACN2B* (log2 FC = −2.1; P = 8.7×10^−4^ at FDR 1.6%) and *GRM3* (log2 FC = 3.3; P = 2.5×10^−3^ at FDR 3.0%).

**Figure 1.**
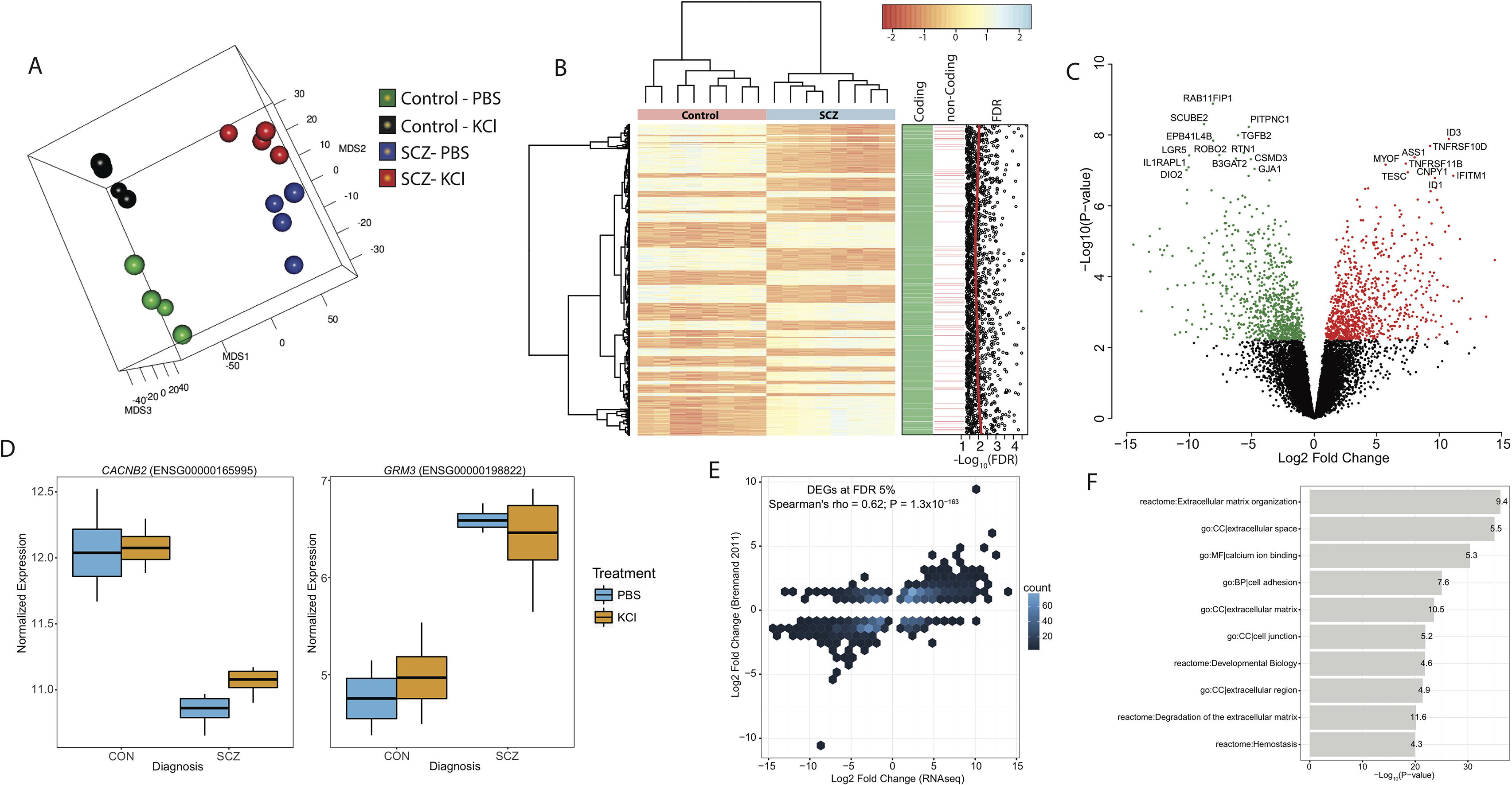
Differential expression between control and SCZ human induced pluripotent stem cell (hiPSC) neurons. **(A)** Multidimensional scaling of RNA sequencing gene expression of hiPSC neurons from each of four control and four cases with SCZ with (KCl) and without (PBS) potassium chloride treatment segregates samples along the three leading fold change dimensions. **(B)** Bivariate clustering of samples (columns) and the 1,669 DEGs for diagnosis at FDR ≤ 0.05 (rows) shows the case-control differences, as marked by the cyan-pink horizontal colorbar at top. Higher or lower expression per sample is marked in cyan or red, respectively. The vertical colorbar indicates whether genes are coding or non-coding and the distribution of FDR for each comparison. **(C)** Volcano plots of −Log10 P value vs. Log2 fold-change for control and SCZ hiPSC neurons. Among the 1,669 DEGs, 854 were up-regulated and 815 were down-regulated in SCZ. The most significant DEGs are indicated. **(D)** Boxplots showing the distributions of normalized gene expression in SCZ vs. controls stratified by treatment with PBS and potassium chloride for *CACNB2* and *GRM3* genes. **(E)** Binned density scatter plot comparing the Log2 fold-change for control vs. SCZ differential expression between the current dataset and a previous hiPSC neuron cohort assayed on microarrays. **(F)** Pathway enrichment analysis across a group of gene sets for known molecular pathways and biological processes (see text for details). The x-axis represents −Log10 P value, and the y-axis denotes pathways. The numbers on each bar indicates the fold enrichment.

We examined differential expression in a previous sample, which generated gene expression data using microarrays from 4 SCZ patients (12 microarrays) and 4 controls (12 microarrays)^9^. While these arrays differ from RNAseq in their capture features, the correlation of test statistics for differential expression in this current dataset compared to the previous was significant: Spearman’s correlation rho = 0.62 (P = 5.7×10^−161^) for the subset of significantly DEGs also present in the previous dataset (n = 1,509 genes) (**Figure 1E**). Pathway enrichment analysis across a broad set of pathways was conducted for interpretation of the list of DEGs. The most enriched categories included pathways related to organization of extracellular matrix, cell adhesion and binding of calcium ion (**Figure 1F**).

### Activity-dependent differential gene expression analysis

We next examined transcriptional changes related to PCID. We identified 1,199 DEGs at FDR ≤ 0.05 (**eTable 3 in the Supplement**). Among the 1,199 genes, 595 were up-regulated and 604 were down-regulated in response to PCID, with a moderate effect (average log2 fold change 1.26; range 0.45-4.82) (**Figure 2A**). Unsupervised clustering shows a clear separation of PCID signatures in controls but less clear in SCZ, indicating a less significant impact of PCID on gene expression in SCZ (**Figure 2B**). **Figure 2C** shows two illustrative examples for *SLC38A2* (log2 FC = 4.2; P = 1.4×10^−8^ at FDR 0.007%) and *THNSL1* (log2 FC = −2.7; P = 2.1×10^−7^ at FDR 0.02%) that are up-and down-regulated in response to PCID, respectively.

**Figure 2.**
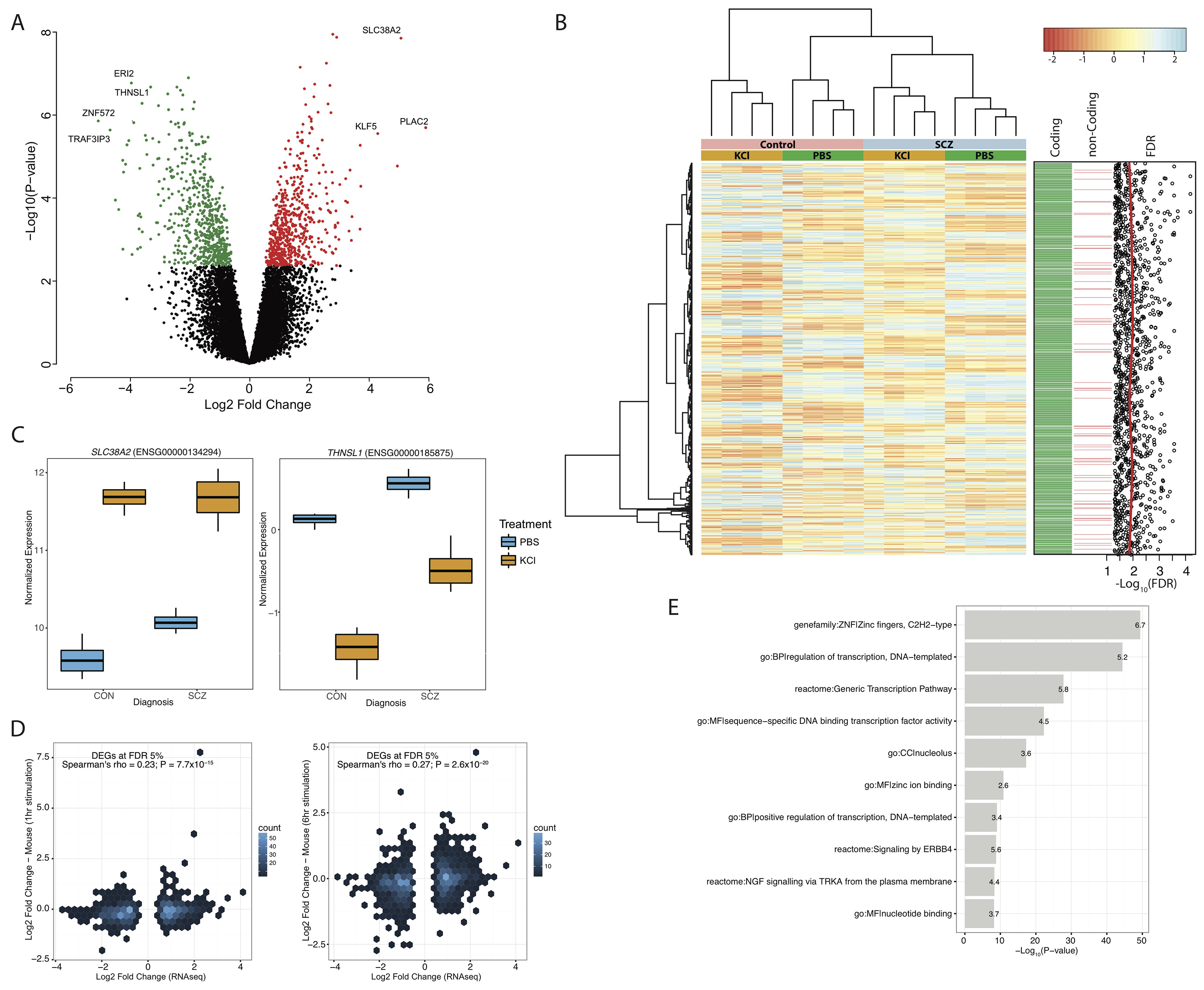
Differential expression between untreated and treated potassium chloride-induce depolarization. **(A)** Volcano plots of −Log10 P value vs. Log2 fold-change for untreated and treated potassium chloride-induce depolarization (PCID). Among the 1,199 genes, 595 were up-regulated and 604 were down-regulated in response to PCID. The most significant DEGs are indicated. **(B)** Bivariate clustering of samples (columns) and the 1,199 DEGs for PCID at FDR ≤ 0.05 (rows) shows the untreated and treated differences in SCZ and controls, as marked by the cyan-pink and orange-green horizontal colorbars at top for diagnosis and PCID treatment, respectively. Higher or lower expression per sample is marked in cyan or red, respectively. The vertical colorbar indicates whether genes are coding or non-coding and the distribution of FDR for each comparison. **(C)** Boxplots showing the distributions of normalized gene expression in SCZ vs. controls stratified by treatment with PBS and potassium chloride for *SLC38A2* and *THNSL1* genes. **(D)** Binned density scatter plots comparing the Log2 fold-change for PCID treatment differential expression between the current dataset and a previous gene expression dataset in mouse cortical neurons after PCID for 1 hour (left plot) or 6 hours (right plot). **(E)** Pathway enrichment analysis across a group of gene sets for known molecular pathways and biological processes (see text for details). The x-axis represents −Log10 P value, and the y-axis denotes pathways. The numbers on each bar indicates the fold enrichment.

We examined PCID signatures in a previous sample, which generated gene expression data in mouse cortical neurons^26^. We found a moderate and significant correlation between our PCID signatures and mouse cortical neuron signatures after PCID for 1 hour (Spearman’s correlation rho = 0.23; P = 7.7×10^−15^) or 6 hours (Spearman’s correlation rho = 0.27; P = 2.6×10^−20^) (**Figure 2D**). Pathway enrichment analysis showed enrichment for molecular functions and biological processes related to transcription factor activity, regulation of transcription and signaling by *ERBB4* and *NRG* (**Figure 2E**).

### PCID signatures are attenuated in SCZ

Based on the more prominent separation of PCID signatures in controls compared to SCZ (**Figure 2B**), we performed separate DEG analysis in each group. Remarkably, whereas 594 genes were PCID DEGs in controls at FDR ≤ 0.05 (**eTable 4 in the Supplement**), only 59 genes were differentially expressed in SCZ (**eTable 5 in the Supplement**). This was not simply an issue of statistical thresholds, as relaxing the statistical criteria for differential expression (FDR ≤ 0.1) identified 1, 112 differentially expressed genes in controls, and only 169 in SCZ, confirming the large difference observed in PCID signatures. We note that PCID signatures were consistent in terms of directionality (up-or down-regulated) in both controls and SCZ (**Figure 3A**).

**Figure 3.**
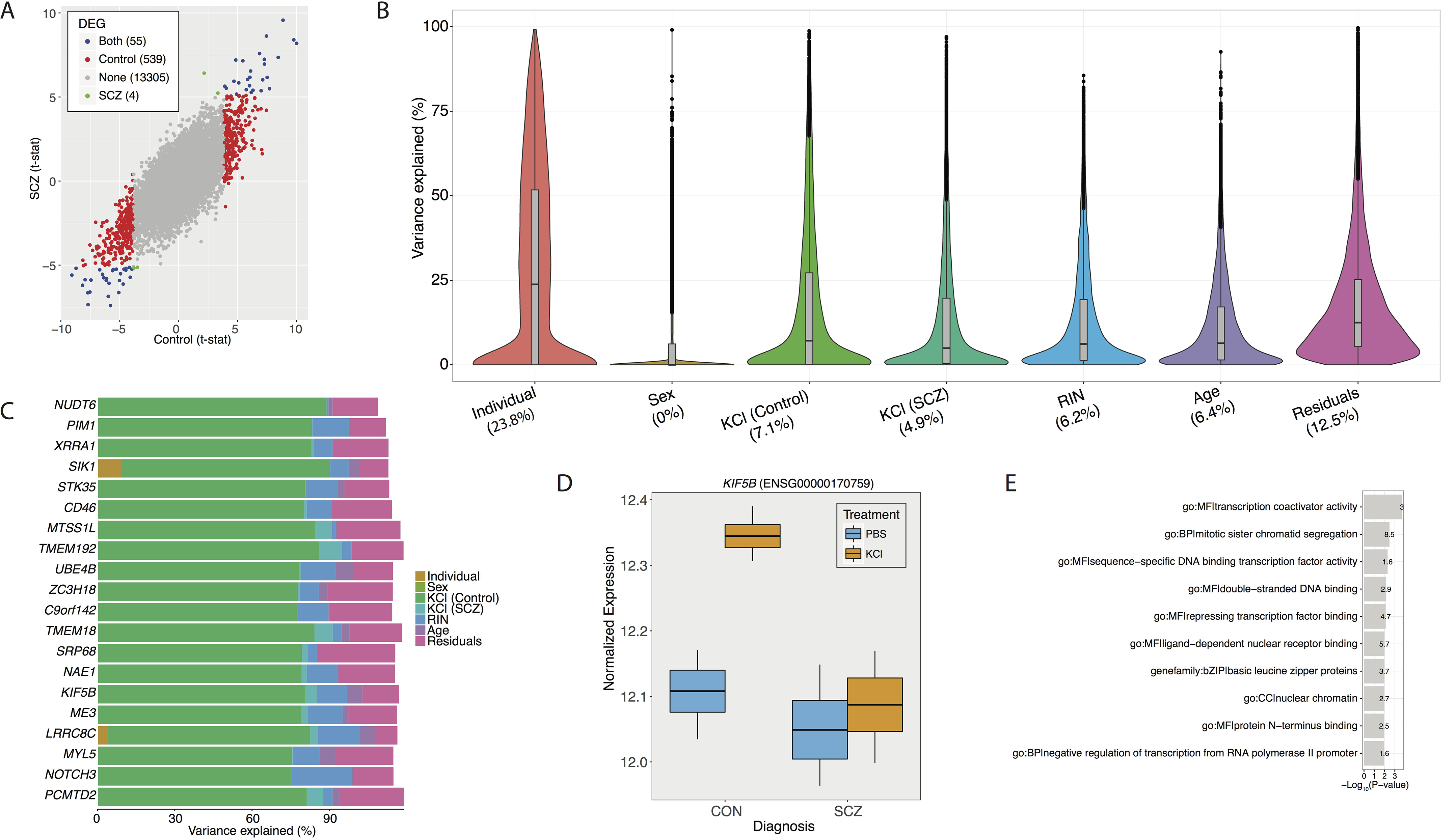
Comparison of gene expression changes after potassium chloride-induce depolarization in controls and SCZ samples. **(A)** Scatter plot comparing the t-statistic for PCID treatment differential expression between controls and cases with SCZ. The numbers in the legend indicate the count of DEGs at FDR ≤ 0.05 for controls only, SCZ only, both SCZ and controls or non-significant genes. **(B)** Violin plots of the percentage of variance explained by individual, PCID in controls and SCZ, sex, age and RIN across all genes. Every gene is represented in the violin plot of each variable. Numbers in the x-axis indicate the median of percentage of variance explained by each experimental variable over all the genes. **(C)** The percentage of variance explained by each experimental variable for the top 20 control signatures (largest difference for PCID variance explained in controls vs. SCZ) is illustrated. **(D)** Boxplot showing the distributions of normalized gene expression in SCZ vs. controls stratified by treatment with PBS and potassium chloride for *KIF5B* gene. **(E)** Pathway enrichment analysis across a group of gene sets for known molecular pathways and biological processes (see text for details). The x-axis represents −Log10 P value, and the y-axis denotes pathways. The numbers on each bar indicates the fold enrichment.

To further quantify the disease effect (control and SCZ) on the PCID transcriptional variability, we applied a linear mixed model for each gene and quantified the total variance of PCID within each diagnostic group^27^. Genome-wide variation across PCID accounted for a median of 7.1% variation in controls; In SCZ the variation explained was 4.9%, confirming our previous observation of attenuated effect of PCID on gene expression in SCZ compared to controls (**Figure 3B**). We then ranked genes based on the difference of the variance of PCID signature in controls versus SCZ. We identified 2,040 genes (control signatures) that over 20% and less of 20% of their variation was explained by treatment in controls and SCZ, respectively. On the other hand, only 1,111 genes (SCZ signatures) had less of 20% and over 20% of their variation explained by treatment in controls and SCZ, respectively. The top 20 control signatures and an illustrative example for *KIF5B* gene are provided (**Figure 3C** and 3D). Pathway enrichment analysis on the top 5% of control signatures showed enrichment for molecular functions and biological processes related to transcription factor activity and regulation of transcription (**Figure 3E**).

### Network analysis identifies Diagnosis and PCID-dependent expression changes

To identify discrete groups of co-expressed genes showing transcriptional differences relevant to diagnosis or PCID, we constructed a co-expression network using the entire data set. Genes were clustered into 15 co-expression modules based on high similarity of expression patterns across samples (**eTable 6in the Supplement**). Genes that clustered into specific modules based on similar co-expression patterns are also DEGs for diagnosis or PCID status, indicating that they participate in common biological processes (**Figure 4A**).

**Figure 4.**
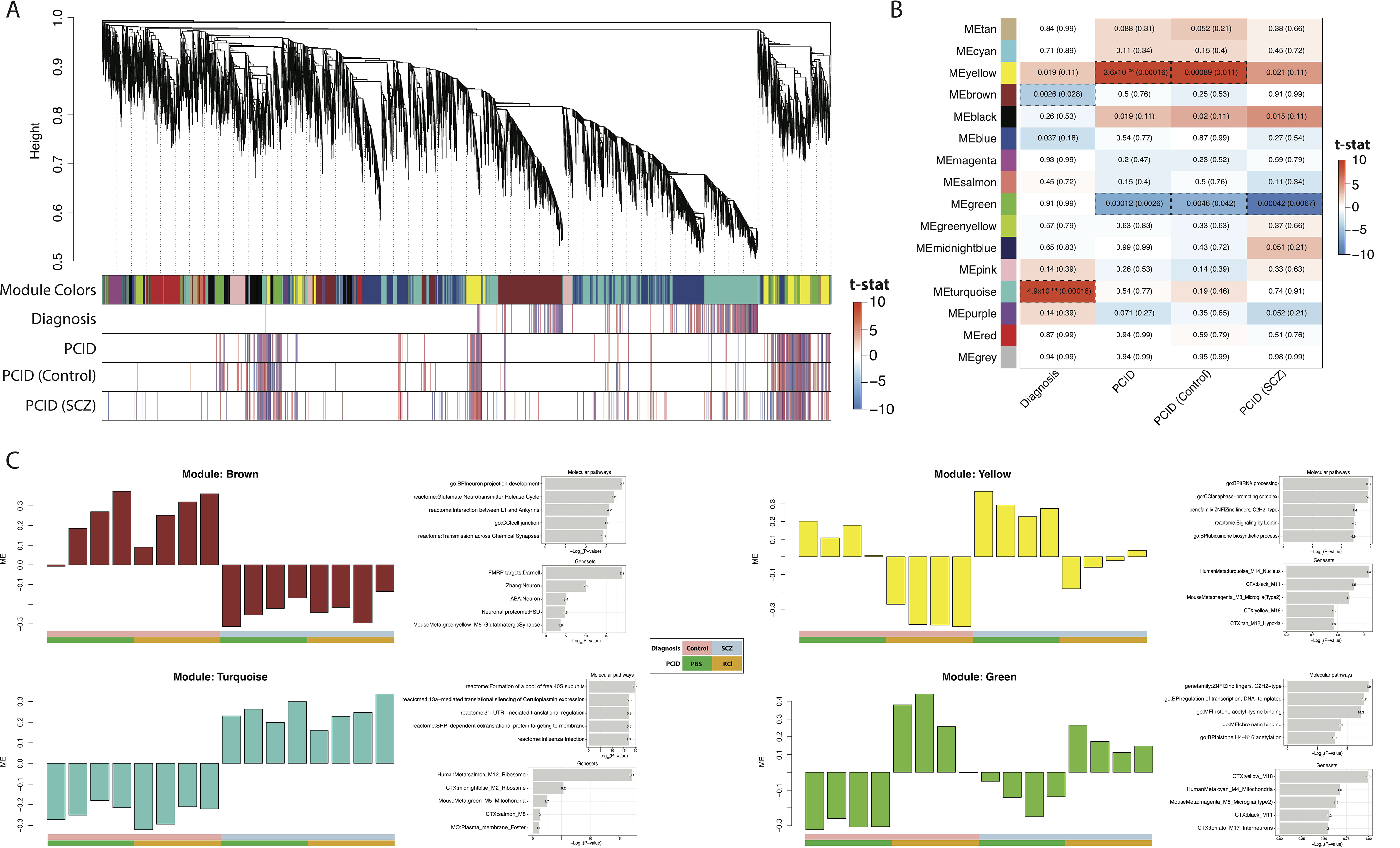
Co-expression networks in potassium chloride-induce depolarization in controls and SCZ samples. **(A)** Hierarchical clustering of genes based on gene co-expression pattern across controls and SCZ cases, with or without PCID. Co-expression modules are represented by color classifiers (Module Colors), noted in the first row of the horizontal colorbar. The “Diagnosis”, “PCID”, “PCID (Control)” and “PCID (SCZ)” colorbars represent the t-statistic from the differential expression analysis. Red indicates up-regulation, while blue indicates down-regulation. **(B)** Heatmap of module eigengenes (ME) association with diagnosis and PCID status. The left panel shows fifteen modules and the gray module (which includes genes that are not co-expressed). The right panel is a color scale for module trait association based on the t-statistic for each experimental variable. Higher or lower expression of each ME per experimental variable is marked in red or blue, respectively. For each association we present the P values and FDR (in parenthesis). Significant associations at FDR < 0.05 are emphasized by box with gapped line. **(C)** Module eigengene patterns and enrichment scores of the top five enriched functional genesets for modules that are significant at FDR ≤ 0.05 with diagnosis (brown and turquoise) or PCID (yellow and green). Left: barplots indicate the level of MEs across samples. Samples are ordered by control-PBS, control-KCl, SCZ-PBS and SCZ-KCl, as illustrated by the key at the bottom. Right: Pathway enrichment analysis across group of gene sets for known molecular pathways and biological processes (top) or hypothesis driven genesets (bottom). The x-axis represents −Log10 P value, and the y-axis denotes pathways. The numbers on each bar indicates the fold enrichment.

We then examined whether the identified co-expression networks recapitulate molecular processes related to diagnosis and PCID status. We used the module eigengene (ME, which is the first principal component of the expression pattern of the corresponding module) to summarize gene expression trajectories across samples, and evaluated the relationship of the 15 module eigengenes with diagnosis and PCID status (**Figure 4B; eTable 7 in the Supplement**). We found 2 modules strongly associated with diagnosis status at FDR ≤ 0.05; one module was up-regulated in SCZ (turquoise) and was enriched for ribosome markers, as well as for genes belonging to 3’-UTR-mediated translational regulation (**Figure 4C**). The other module was down-regulated in SCZ (brown) and was enriched for neuronal markers, as well as for genes belonging to FMRP targets (**Figure 4C**).

We also found 2 modules strongly associated with PCID status at FDR ≤ 0.05; one module was up-regulated with KCl treatment (yellow) and was enriched for genes belonging to tRNA processing and ZNF gene family (**Figure 4C**). The other module was down-regulated with KCl treatment (green) and was enriched for genes belonging to regulation of transcription and ZNF gene family (**Figure 4C**). Both modules were significant when we examined the association of ME with PCID in control samples, while only the green module was significant in SCZ. These results are consistent with the less significant effect of PCID on gene expression (and MEs) in SCZ compared to controls.

### Genetic variants associated with SCZ are enriched in differentially expressed genes

To further interpret the co-expressed modules, we examined whether modules are enriched for genes that have been previously associated with SCZ and other neurodevelopmental illnesses (autism, epilepsy and intellectual disability) (**Figure 5A; eTable 8 in the Supplement**). Common SCZ risk variants defined based on the most recent large-scale GWAS from PGC2^1^, were enriched for 5 modules at P < 0.05; interestingly 3 out of the 5 modules were associated with disease status -- brown (P = 0.003) and turquoise (P = 0.016) -- or PCID -- yellow (P = 0.032). A significant overlap of the yellow module (P=0.048) was identified with genes that lie within genomic regions, which have been associated with large cytogenomic deletions in SCZ; no significant effect was observed for SCZ-associated duplicated regions.

**Figure 5.**
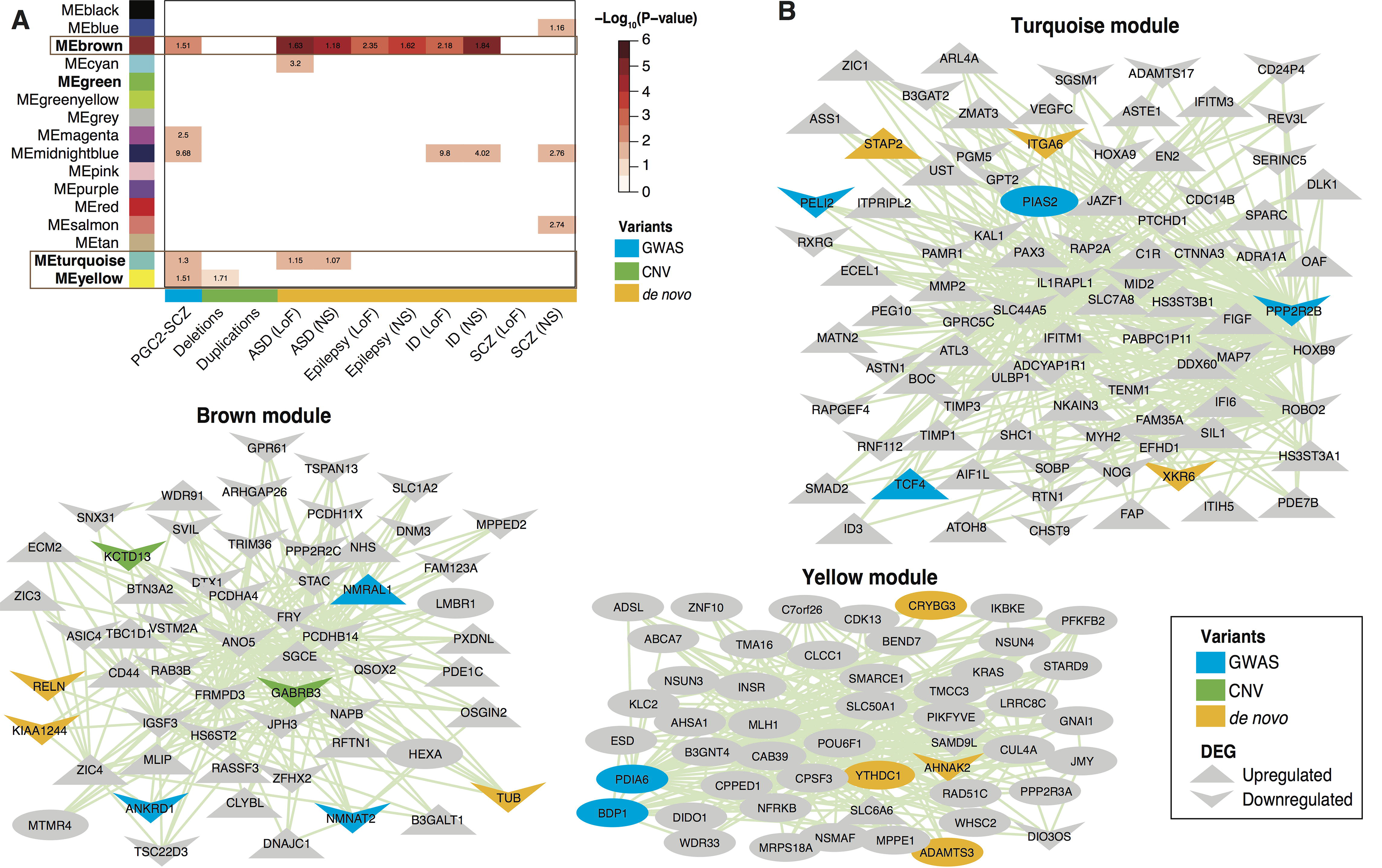
Enrichment of co-expression modules with genetic risk variants for SCZ and other neurodevelopmental diseases. **(A)** Heatmap of enrichment of modules with genetic risk variants for SCZ -- common variants from PGC2-SCZ analysis, copy number variants or *de novo* mutations -- and *de novo* mutations across neurodevelopmental diseases, including autistic spectrum disorder (ASD), intellectual disability (ID) and epilepsy. The left panel shows fifteen modules and the gray module (which includes genes that are not co-expressed). The right panel is a color scale for enrichment based on the −Log10 P value for each module with genetic variants. More significant enrichment is marked in red. For each association we present the ratio of observed vs. expected overlapping genes. Only significant enrichments with P ≤ 0.05 are shown. In bold fonts and within box are modules that are significantly associated with disease and PCID and have a significant enrichment for genetic variants. **(B)** Intramodular hub genes for modules that are significantly associated with disease and PCID and have a significant enrichment for genetic variants are shown. Genes that have a genetic association for SCZ (GWAS, CNV or *de novo*) or are DEGs for diagnosis (brown and tuqrquoise) or PCID (yellow) are illustrated. NS: non-synonymous; LoF: loss-of-function mutations.

The brown module was significantly enriched for nonsynonymous and loss-of-function *de novo* mutations across multiple neurodevelopmental diseases, including autism, epilepsy and intellectual disability (ID), but not SCZ. In addition, the turquoise module was significant enriched for *de novo* mutations in autism. SCZ *de novo* mutations were enriched for 3 modules; none of these modules were associated with disease or PCID status. Finally, we examined the effect of SCZ-associated genetic signal in aggregate and identified a significant effect for 3 out of the 4 modules that were associated with disease or PCID status: brown (Fisher’s P = 0.002), turquoise (Fisher’s P = 0.044) and yellow (Fisher’s P = 0.018). The hub nodes (most connected genes) are illustrated for the brown, turquoise and yellow modules (**Figure 5B**).

## DISCUSSION

Neuronal depolarization regulates synaptic function by inducing gene expression and protein synthesis of synaptic molecules in dendrites^8^. This study shows that hiPSC neurons display activity-dependent changes on gene expression, when treated with potassium chloride, similar to activity-dependent changes in neurotransmitter release that have previously been reported in neurons differentiated from these same hiPSC lines^28^. These genes participate in molecular functions and biological processes related to regulation of transcription and they show consistent changes with gene expression studies in mouse cortical neurons^26^. Thus, the transcriptome of hiPSC neurons is induced in response to neuronal depolarization, indicating that they are a relevant platform for the study of activity-dependent changes in neuropsychiatric diseases.

Compared to controls, hiPSC neurons derived from cases with SCZ showed alterations of gene expression consistent with a previous study^9^ and are enriched in molecular pathways related to organization of extracellular matrix and cell adhesion. These extracellular matrix proteins include gene families such as neural cell adhesion molecules, neuroligins and neurexins that are critical for cellular differentiation and migration^29^, and are postulated as a possible mechanism for the etiopathogenesis of SCZ^30^. Previous studies have demonstrated that SCZ NPCs have abnormal expression for extracellular matrix and cell adhesion genes^10,31^, accompanied by deficits in migration^10^.

By examining the higher order organization of the transcriptome, we identified specific modules to be associated with disease status and PCID. We identified two modules that were significantly deregulated in SCZ and two modules that are altered in response to PCID. Similarly, to differential expression analysis, we observed a blunted effect of activity-dependent changes of module activity in SCZ. By testing the enrichment of modules with SCZ candidate genes, we found a strong enrichment of genetic variants with the disease-associated modules and the PCID-associated network that was only activated in controls, but not SCZ. These data support the notion that the impairment of different but not mutually exclusive molecular pathways contributes to the etiology of SCZ. For instance, SCZ may arise from early defects in neurodevelopment, followed by global dysregulation of synaptic function and impairment of neurotransmission. This is consistent with findings in the brown module that participates in neurodevelopment and synaptic function, has a strong enrichment for SCZ variants and is associated with the disease status. Alternatively, another possibility is that causative factors involve dysregulation of activity-dependent signalling pathways either locally at the synapse or globally, leading to deficits in activity-dependent gene transcription. This is consistent with results in the yellow module, which is enriched for SCZ risk variants and has a strong association with PCID in controls.

While our study shows convergence of genetic variants to specific dysregulated co-expressed networks, it fails to identify driver genes. This can be the focus of future studies, where incorporating larger sample sizes and including additional time-points for PCID treatment would allow to generate directed networks, such as causal probabilistic networks, and identify specific genes that drive abnormalities in SCZ. Finally, future studies should collect additional functional data, such as quantification of secreted neurotransmitters and electrophysiological measurements, which would allow investigating the effect of dysregulated gene expression on neurons’ physiology.

In conclusion, here we examined the effect of PCID using hiPSC neurons from cases with SCZ and controls and identified co-expressed networks that are dysregulated in SCZ and in response to PCID. We found that these networks are enriched for SCZ and other neurodevelopmental diseases risk variants, providing an additional functional dataset that can provide further insight into the function and mechanism of SCZ candidate genes. Finally, we show that activity dependent changes of gene expression are relevant to SCZ etiology and hiPSC neurons can be utilized to further examine the effect of neuronal activity on the SCZ transcriptome in future studies.

## Author contributions

Drs Roussos and Brennand had full access to all of the data in the study and take responsibility for the integrity of the data and the accuracy of the data analysis.

*Study concept and design:* Roussos, Barry, Brennand.

*Generation of cells:* Brennand. *Generation of sequencing data:* Kaczorowski, Barry.

*Statistical analysis:* Roussos, Guennewig.

*Drafting of the manuscript:* Roussos, Barry, Brennand.

*Obtained funding:* Roussos, Barry, Brennand.

## Conflict of Interest Disclosures

None reported.

## Funding/Support

This work was supported by the National Institutes of Health (R01AG050986 Roussos, R01MH101454 Brennand and R01MH106056 Brennand), New York Stem Cell Foundation (Brennand), Brain Behavior Research Foundation (Roussos and Brennand), Alzheimer’s Association (NIRG-340998 Roussos), the Veterans Affairs (Merit grant BX002395 Roussos), the Swiss National Science Foundation (P2EZP3_152143 Guennewig) and a Geoff and Dawn Dixon fellowship (Guennewig).

## Role of the Funder/Sponsor

The funders had no role in the design and conduct of the study; collection, management, analysis, and interpretation of the data; preparation, review, or approval of the manuscript; and decision to submit the manuscript for publication.

